# In vitro measurements of protein–protein interactions show that antibody affinity governs the inhibition of SARS-CoV-2 spike/ACE2 binding in convalescent serum

**DOI:** 10.1101/2020.12.20.422820

**Authors:** Sebastian Fiedler, Monika A. Piziorska, Viola Denninger, Alexey S. Morgunov, Alison Ilsley, Anisa Y. Malik, Matthias M. Schneider, Sean R. A. Devenish, Georg Meisl, Adriano Aguzzi, Heike Fiegler, Tuomas P. J. Knowles

## Abstract

The humoral immune response plays a key role in suppressing the pathogenesis of SARS-CoV-2. The molecular determinants underlying the neutralization of the virus remain, however, incompletely understood. Here, we show that the ability of antibodies to disrupt the binding of the viral spike protein to the angiotensin-converting enzyme 2 (ACE2) receptor on the cell, the key molecular event initiating SARS-CoV-2 entry into host cells, is controlled by the affinity of these antibodies to the viral antigen. By using microfluidic antibody-affinity profiling, we were able to quantify the serum-antibody mediated inhibition of ACE2–spike binding in two SARS-CoV-2 seropositive individuals. Measurements to determine the affinity, concentration, and neutralization potential of antibodies were performed directly in human serum. Using this approach, we demonstrate that the level of inhibition in both samples can be quantitatively described using the binding energies of the binary interactions between the ACE2 receptor and the spike protein, and the spike protein and the neutralizing antibody. These experiments represent a new type of in-solution receptor binding competition assay, which has further potential areas of application ranging from decisions on donor selection for convalescent plasma therapy, to identification of lead candidates in therapeutic antibody development, and vaccine development.

## Introduction

The COVID-19 pandemic is not only causing a major public health crisis but also unprecedented economic challenges. One of the features of COVID-19 is the variability of disease outcomes, with a large majority of patients presenting mild or no symptoms, while others may become severely ill or die^1^. Independently of the disease trajectory, the majority of acute respiratory syndrome coronavirus 2 (SARS-CoV-2) infections trigger a response of the adaptive immune system^2–4^. Essential for this immune response are antibodies that neutralize SARS-CoV-2 (NAbs) by preventing virus binding to the host-cell receptor angiotensin-converting enzyme 2 (ACE2)^5–13^. Effective NAbs target the receptor binding domain (RBD) of viral spike proteins and hence inhibit ACE2 binding^13^. Such antibodies and their characterization are of great interest either for therapeutic approaches such as plasmapheresis^14^ or as structural templates for rational vaccine design^15^.

Consequently, there is a need for assays that identify the most potent NAbs reliably and efficiently. The gold standard to detect NAbs is a virus-neutralization test (VNT) on live cells which can either be performed with live viruses or pseudo-viruses^16,17^. While these assays are well established, VNTs are time-consuming and logistically challenging, require several days to obtain results, and necessitate stringent biosafety measures to be in place for handling the live virus. Furthermore, it is challenging to standardize VNTs to allow consistency among different laboratories, studies, and patient cohorts^18^. This situation has motivated the search for easier to use and faster alternatives to VNTs in the form of surrogate VNTs (sVNTs). Such assays are typically based on an enzyme-linked immunosorbent assay (ELISA), which provide information on the titers of NAbs in a patient sample^19^. Neither the standard VNTs, nor the ELISA-based sVNTs can, however, provide information on whether virus neutralization is achieved by a high concentration of NAbs with weak binding affinities or a lower concentration of NAbs with tight binding affinities. This information, however, is of particular importance not just for understanding the fundamental physicochemical parameters that underlie neutralization but has also potential applications for selection of suitable donors for plasmapheresis.

Here, we investigate whether NAb efficacy can be predicted based on affinity by assessing the binding energies that drive ACE2 displacement from the SARS-CoV-2 RBD. Using microfluidic antibody-affinity profiling (MAAP), we demonstrate that a ternary equilibrium of ACE2, SARS-CoV-2 spike S1 (S1), and antibodies from seropositive, recovered individuals predicts and quantifies the inhibition of ACE2–S1 binding. MAAP is an in-solution assay that determines both the effective dissociation constants (*K*_D_) of polyclonal serum antibodies and their concentrations^21,22^. Based on these findings, we suggest that the ratio of the dissociation constants of ACE2–S1 binding and NAb–S1 binding (r = *K*_D,ACE2_ / *K*_D,NAb_) together with the experimentally determined concentration of NAbs in serum are excellent measures to predict the inhibitory potential of the polyclonal antibody response. Such quantitative information on antibody-mediated inhibition of ACE2–S1 binding is crucial to drive decision making in donor selection for plasmapheresis, lead identification in the development of therapeutic antibodies, and in vaccine development.

## Results and Discussion

### Affinity measurements in pre-pandemic control serum using recombinant ACE2, S1, and a monoclonal neutralizing anti-S1 antibody

In this study, we investigate whether the affinity of anti-S1 antibodies present in human serum samples is a predictor for their neutralization efficacy. For this purpose, we established a novel assay using microfluidic diffusional sizing (MDS, **Figure 1A**) utilizing a recombinant, monoclonal NAb (NAb1) spiked into SARS-CoV-2 pre-pandemic control serum in the presence of ACE2 and S1. The NAb-mediated inhibition of ACE2–S1 binding is physically determined by the ternary equilibrium ACE2–S1–NAb, which is described quantitatively by the *K*_D_ values of the two binary equilibria, ACE2–S1 and NAb–S1 (refer to **Materials and Methods** for details). Thus, to validate the data generated on the ternary system, we determined the *K*_D_ values of the two underlying binary reactions.

**Figure 1.**
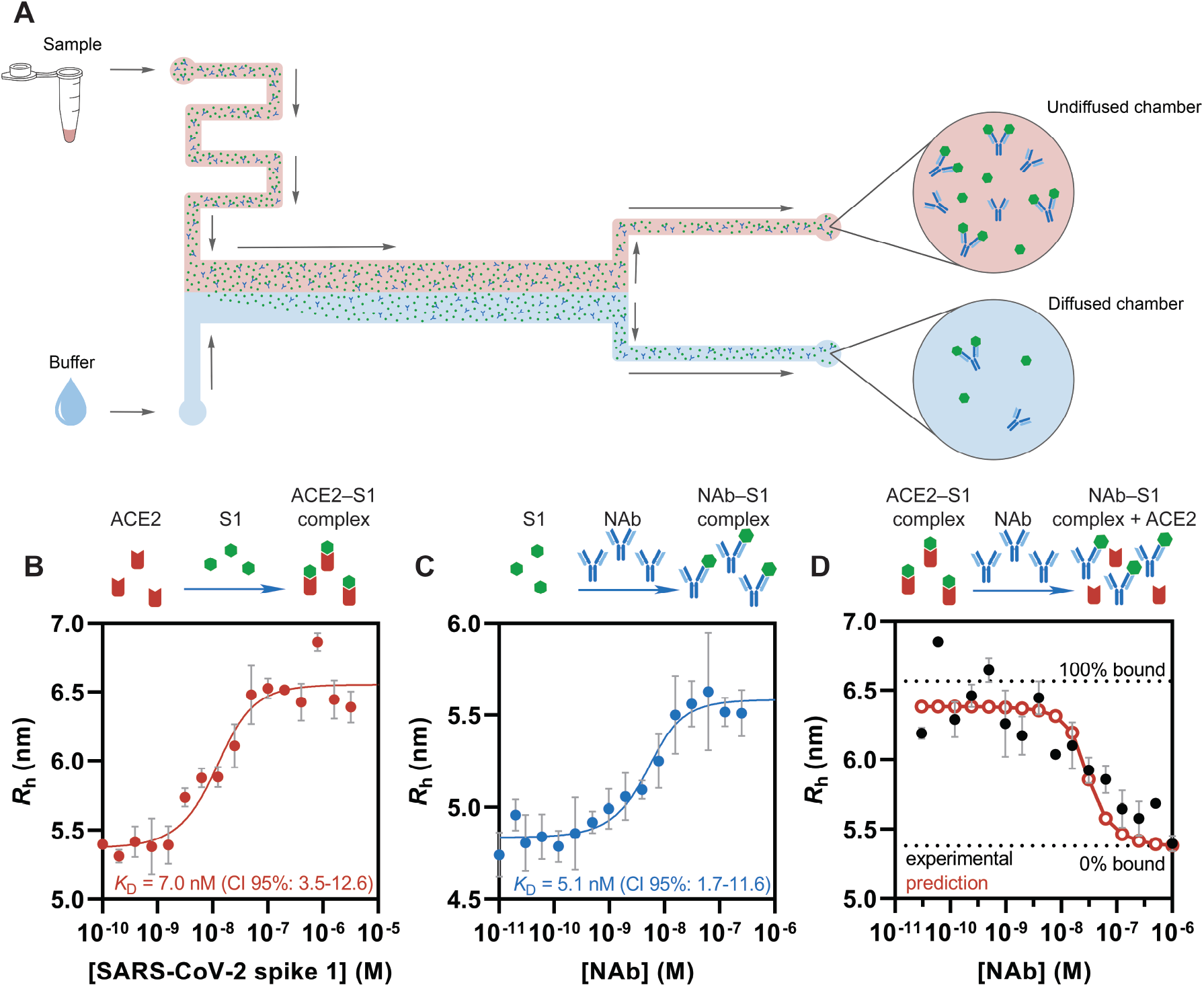
Measurement of S1 affinities and NAb-mediated inhibition of spike binding in 90% pre-pandemic control serum by microfluidic diffusional sizing (MDS). **A**) On a microfluidic device, a fluorescently labeled species is introduced into a diffusion chamber alongside an auxiliary stream under laminar flow conditions. Migration of the fluorescently labeled species into the auxiliary stream is diffusion controlled at low Reynold’s number, which enables measurement of the hydrodynamic radius, *R*_h_, from the ratio of fluorescence intensities in the diffused chamber and the undiffused chamber. **B**) and **C**) Equilibrium binding of ACE2–S1 and NAb1–S1, respectively, measured by MDS in the presence of 90% pre-pandemic control serum. Lines indicate fits in terms of a binary equilibrium (**Equation 2**). Errors for *K*_D_ are 95% confidence intervals. Error bars represent the standard deviation from triplicate measurements. **D**) Inhibition of ACE2–S1 binding by a recombinant, monoclonal NAb (NAb1) in 90% pre-pandemic control serum measured by MDS. Fluorescently labeled ACE2 at 10 nM and S1 at 50 nM (predicted 80% of ACE2 bound to S1) were mixed with increasing concentrations of NAb1. NAb1 displaces labeled ACE2 from S1, causing *R*_h_ to decrease to the level of unbound ACE2. Red, empty circles indicate predicted change in *R*_h_ in terms of a ternary equilibrium (**Equation 11**) based on the underlying *K*_D_ values of ACE2–S1 and of NAb1–S1. Dotted lines indicate *R*_h_ of free ACE2 and ACE2–S1 complex taken from Figure 1B. Error bars represent the standard deviation from triplicate measurements.

First, we measured the *K*_D_ of ACE2–S1 binding in the presence of 90% pre-pandemic control serum (**Figure 1B**). As the S1 concentration was increased, more of the fluorescently labeled ACE2 was found in the protein complex and the effective hydrodynamic radius (*R*_h_) increased. Based on a 1:1 equilibrium binding model (**Equation 2**), the *K*_D_ of ACE2–S1 was determined to be 7.0 nM (95% CI: 3.5–12.6). Next, we measured S1 binding to NAb1 in 90% pre-pandemic control serum (**Figure 1C**). To do so, we observed the increase in *R*_h_ caused by complex formation of fluorescently labeled S1 and NAb1. The resulting *K*_D_ of NAb1–S1 was determined to be 5.1 nM (95% CI: 1.7–11.6).

### Inhibition of ACE2–S1 binding in pre-pandemic control serum by a monoclonal neutralizing anti-S1 antibody

After having characterized the binding between ACE2 and S1 as well as between NAb1 and S1, we investigated whether NAb1-mediated inhibition of ACE2–S1 binding occurs in the ternary mixture of ACE2, S1, and NAb1. The addition of NAb1 to a mixture of fluorescently labeled ACE2 at a concentration of 10 nM and S1 at a concentration of 50 nM resulted in a reduction of *R*_h_, as NAb1 bound to S1, thereby displacing ACE2 from the complex (**Figure 1D**). At NAb1 concentrations of approximately 1 µM, the effective *R*_h_ of the ACE2–S1–NAb1 mixture corresponded to *R*_h_ of unbound ACE2, indicating complete inhibition of ACE2–S1 binding. Quantitatively, NAb1-mediated inhibition is controlled by the *K*_D_ values of the two binary interactions, ACE2–S1 and NAb1–S1, as well as by the total concentrations of the three interaction partners, as outlined in **Materials and Methods**. For the individual binary reactions, the *K*_D_ values were *K*_D_ = 7.0 nM (CI 95%: 3.5–12.6) for ACE2–S1 and *K*_D_ = 5.1 nM (CI 95%: 1.7–11.6) for NAb1–S1 (**Figures 1B** and **C**). These two affinities enabled us to predict the inhibition isotherm based on a competition binding model (**Equation 11**)^23^. In the ternary mixture used for the competition experiment ([ACE2]_0_ = 10 nM, [S1]_0_ = 50 nM, and 30 pM ≤ [NAb1]_0_ ≤ 1 µM), 80% of ACE2 is predicted to be bound to S1 in the absence of NAb1, and [NAb1]_0_ ≥ 500 nM should completely displace ACE2 from S1 (**Figure 1D, red**). The experimental inhibition isotherm was in remarkably good agreement with the prediction, demonstrating that NAb1 targets the spike RBD and competes with ACE2 for S1 binding, and that the binary interactions sufficiently describe the ternary inhibition interaction.

For a quantitative analysis of these inhibition data, we fitted globally both the two binary binding isotherms (taken from **Figures 1B** and **C**) and the ternary inhibition isotherm (taken from **Figure 1D**).

The global fit utilizes the *K*_D_ values of ACE2–S1 and NAb1–S1 as well as *R*_h_ of unbound ACE2 as global fitting parameters (**Figure 2A**). The *K*_D_ values obtained through the global fit are in agreement with those extracted from the individual binary fits, demonstrating that NAb1-mediated inhibition is driven by the binding energies of the competing protein complexes ACE2–S1 and NAb1–S1. Next, we used *K*_D,ACE2_ and *K*_D,NAb1_ to calculate equilibrium concentrations of free ACE2, S1, and NAb1 as well as of the complexes formed by ACE2–S1 and NAb1–S1.

**Figure 2.**
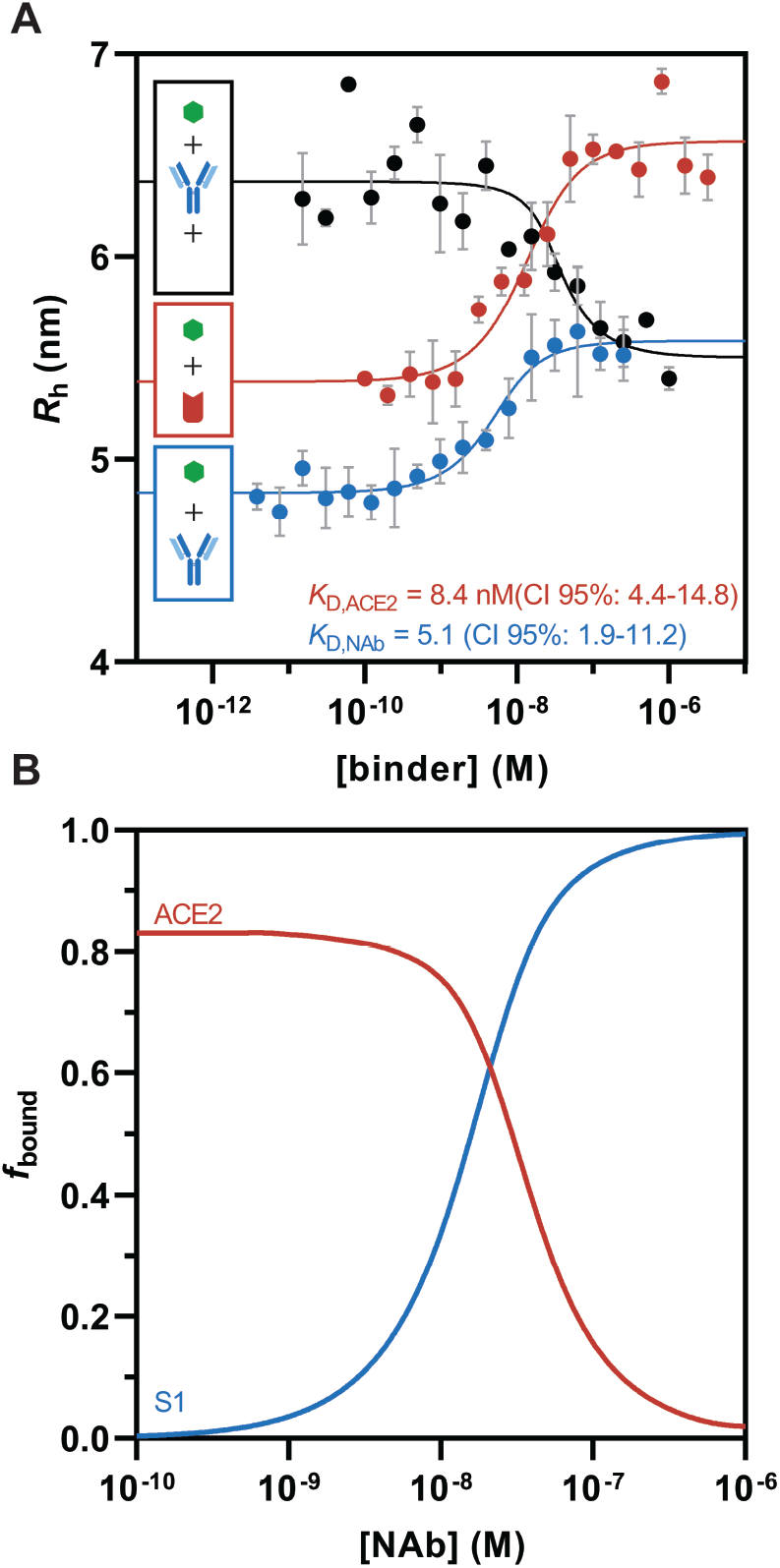
Global analysis of NAb1-induced inhibition of ACE2–S1 binding in 90% pre-pandemic control serum. **A**) Binary binding of ACE2–S1 (red), binary binding of NAb1–S1 (blue), and the ternary equilibrium of labeled ACE2, S1, and NAb1 (black). For analysis, binding and inhibition isotherms were fitted globally in terms of **Equations 2** and **16**, respectively, using *K*_D_ values of ACE2–S1 and NAb1–S1 as well as *R*_h_ of unbound ACE2 as shared parameters. Errors for *K*_D_ are 95% confidence intervals. Error bars are standard deviations of triplicate measurements. **B**) Speciation curves of the ACE2–S1–NAb1 equilibrium as derived from the global fit, displaying fractions of NAb1-bound S1 (blue) and S1-bound ACE2 (red).

Based on those concentrations, we determined the S1-bound fraction of ACE2 and the NAb1-bound fraction of S1 to evaluate NAb1 efficacy (**Figure 2B**).

An effective NAb will prevent viral spike proteins from binding to ACE2, which is key to prevent host-cell infection. *In vivo*, this is accomplished through a combination of NAb–S1 affinity and NAb concentration, as the other parameters that govern the equilibrium, such as ACE2–S1 affinity and the concentrations of ACE2 and S1, cannot be influenced by the host. Thus, the most effective NAbs will have *K*_D_ values considerably lower than the *K*_D_ of ACE2–S1 and concentrations considerably higher than their *K*_D_ for S1 binding. This condition is directly described by the ratio r = *K*_D,ACE2_ / *K*_D,NAb_, which is provided by our assay. For example, at r = 10, the concentration of NAb could be ten times lower than the concentration of ACE2 to still bind the same amount of S1.

### Microfluidic antibody-affinity profiling and neutralization potency in serum of SARS-CoV-2 seropositive individuals

After establishing the assay using a recombinant, monoclonal NAb, we applied the same approach to the quantification of the potency of polyclonal anti-S1 antibody responses in COVID-19 patient samples. To do so, we analyzed blood serum of two individuals who had tested seropositive for anti-SARS-CoV-2 IgG and IgM. Sample 1 was obtained from a 52-year old white female with mild COVID-19 symptoms, while sample 2 was derived from a 44-year old asymptomatic white male.

To determine the *K*_D_ of anti-S1 antibody–S1 binding, we mixed fluorescently labeled S1 at a constant concentration with patient serum at various dilutions and determined *R*_h_ as an indicator of complex formation (**Figures 3A** and **C**). Since the anti-S1 antibody concentration in the two patient sera was unknown, we extracted this information from the equilibrium binding isotherms by using the concentration of antibody-binding sites as a fitting parameter^21,22^. To obtain well-constrained binding-site concentrations, we measured antibody-binding isotherms at three different concentrations of fluorescently labeled S1. To also assess the neutralization potential of the anti-S1 antibodies in the patient sera, we obtained inhibition isotherms from ternary mixtures of fluorescently labeled ACE2, S1 and patient serum at various concentrations (**Figures 3B** and **D**). Both seropositive samples inhibited binding of ACE2 to S1. While sample 1 reached complete inhibition at a serum concentration of 90%, ACE2 was not completely displaced from S1 in sample 2, which could either be due to weak-affinity antibodies or a too low concentration of antibodies.

**Figure 3.**
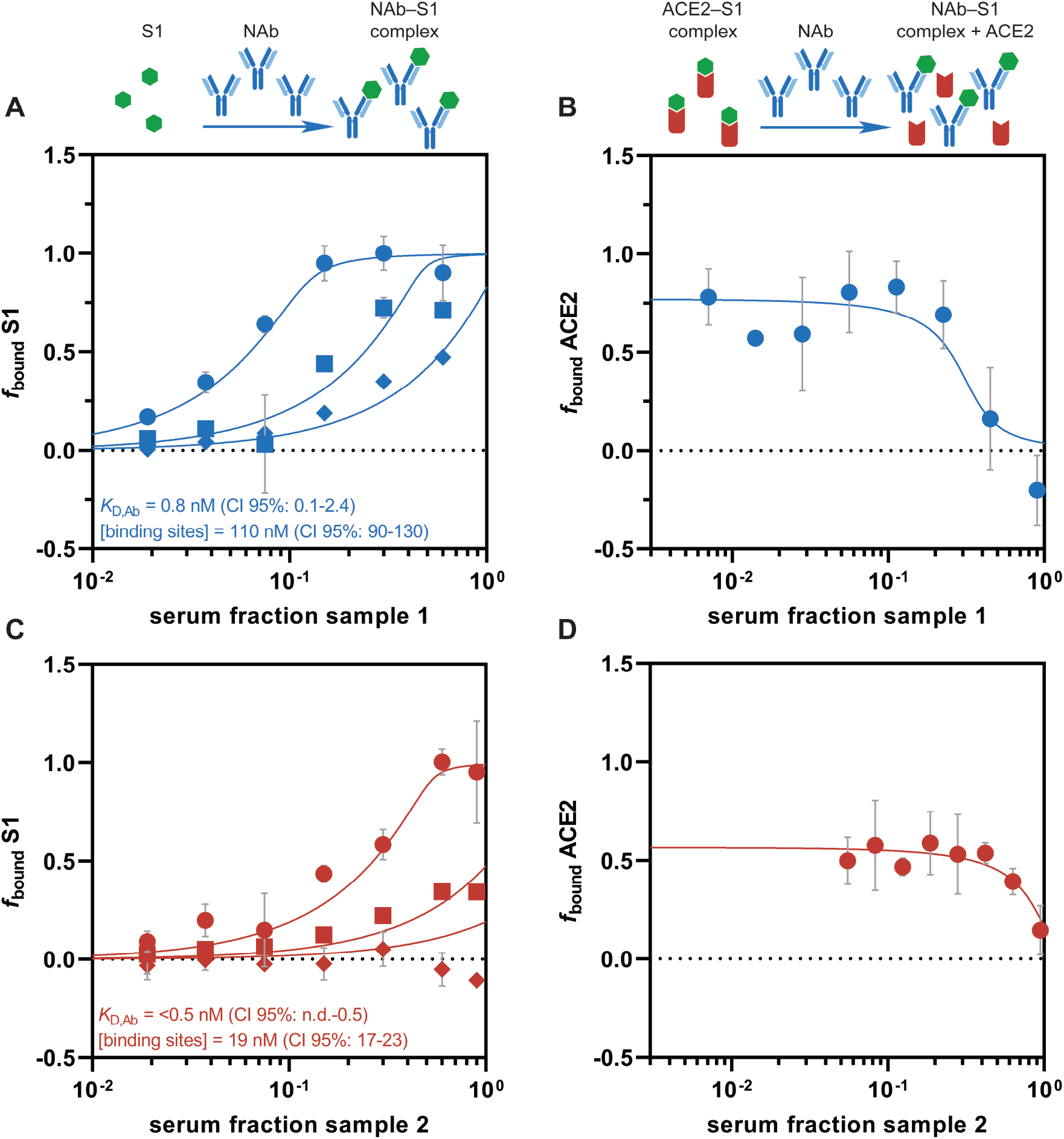
Global fits of antibody–S1 binding and antibody-mediated inhibition of ACE2–S1 binding for SARS-CoV-2 antibody positive, convalescent serum sample 1 (A–B) and sample 2 (C–D). **A**) and **C**) Fluorescently labeled S1 at concentrations of either 10 nM (circles), 40 nM (squares) or 100 nM (diamonds) was mixed with increasing concentrations of patient serum, and binding of antibodies was assessed as an increase in *R*_h_. Fits in terms of **Equation 2. B**) and **D**) Inhibition of ACE2–S1 binding by serum antibodies. Fluorescently labeled ACE2 and S1 were mixed with increasing concentrations of serum. Serum antibodies displace labeled ACE2 from S1, causing *R*_h_ to decrease. Fits in terms of **Equation 16**. Shared values of antibody *K*_D_ were used to describe panels A and B (sample 1) and C and D (sample 2). In all fits, *K*_D,ACE2_ was set to 7.0 nM and used as constant parameter.

For each serum sample, we generated three binary binding isotherms and one inhibition isotherm which we used as a combined set of data to extract *K*_D_ values of antibody–S1 binding, concentrations of antibody-binding sites and neutralization potency. As demonstrated with NAb1 (**Figure 2A**), we globally fitted binary antibody–S1 (**Equation 2**) and ternary antibody-mediated inhibition (**Equation 16**) using the *K*_D_ of antibody–S1 binding as a shared parameter across all sets of data. The *K*_D_ value of ACE2–S1 binding in human serum (7.0 nM) was used as a constant in the fit, based on the data shown in **Figure 1B**. In both patients, global analysis of binding and inhibition resulted in *K*_D_ values in the sub-nanomolar range, which is in line with the values obtained by fitting the binding isotherms without the inhibition isotherm (**Table S1** and **Figure S1**). A MAAP study on a larger number of patient samples showed a similar range of affinities among anti-spike serum antibodies^21^. To investigate the robustness of the *K*_D_ values derived by the global fits, we also analyzed the data using Bayesian inference^21,22^. While the agreement of Bayesian inference with the global fits was excellent, in both patient samples the posterior probability for the *K*_D_ distribution provided strong constraints for the upper bound but weak constraints for the lower bound, thus providing an upper limit on the *K*_D_ value and indicating a tight binding interaction. The best-fit values from the global fits converged to this upper limit representing the higher end of *K*_D_ values that describe the data. For sample 2, the antibody-binding affinity was tighter than for sample 1. The concentration of antibody-binding sites was 110 nM (CI 95%: 90–130) and 19 nM (CI 95%: 17–23) for sample 1 and sample 2, respectively, values which are within the concentration range observed in an earlier MAAP study of a larger number of samples^21^. In both serum samples, the *K*_D_ values of antibody–S1 binding were considerably lower than the concentration of antibody-binding sites, a condition which is required for effective antibody binding.

Moreover, in both samples the ratio r = *K*_D,ACE2_ / *K*_D,NAb_ is 10 or higher, indicating that both individuals produced antibodies against S1 with good neutralization potential. The global fit describes the inhibition isotherms (**Figures 3B** and **D**) of both patient sera remarkably well, demonstrating that antibody-mediated inhibition of the ACE2–S1 interaction is driven by serum-antibody affinity. Furthermore, this indicates that most of the anti-S1 antibodies raised by these two individuals target the RBD of S1 and compete with ACE2 binding as antibodies binding to other epitopes would result in data that deviate from the combined analysis of binary and ternary equilibria used here.

## Conclusions

In this study, we have used microfluidic antibody-affinity profiling to measure independently the binding affinities of the two molecular interactions involved in neutralizing SARS-CoV-2 cell entry, namely that between ACE2–spike and spike–NAb. Furthermore, we have demonstrated that the knowledge of these two interaction energies enables the quantitative description of the ability of patient antibodies to inhibit the binding of the viral spike protein to the host cell ACE2 receptor. The knowledge of both affinity and concentration of anti-S1 antibodies provides valuable information that has the potential to support decision making in research as well as clinical practice. For example, in convalescent plasma therapy a donor with high-affinity, high-concentration anti-RBD antibodies would be most suitable as donor antibodies are diluted c.a. by a factor of ten during the procedure. Specifically, based on this analysis, sample 1 containing high-affinity antibodies at a concentration of 110 nM should be preferred for plasmapheresis over sample 2 which also contains high-affinity antibodies but at a significantly lower concentration of 19 nM. Furthermore, for the selection of autoantibodies as therapeutic candidates it is crucial to be able to deconvolute antibody titers into the component fundamental quantities of affinity and concentration. Indeed, a given antibody titer, as measured for example by conventional ELISA, could arise due to a low concentration of very tightly binding antibodies, which would be attractive candidates for further developments, or by contrast from a high concentration of weakly binding antibodies which are less interesting as leads for further optimization for therapeutic use. Finally, the ability to measure concentration and affinity provides an added level of granularity in understanding the diverse immune responses characteristic of COVID-19 infections and has the potential to evaluate and understand the potency of vaccines, which in order to be optimally effective must generate immune responses that lead to high-affinity virus-neutralizing antibodies at high concentrations.

## Materials and Methods

### Fluorescent labeling of proteins

SARS-CoV-2 spike S1 (S1N-C52H4, ACROBiosystems) and the extracellular domain of ACE2 (AC2-H52H8, ACROBiosystems) were each reconstituted in phosphate buffered saline (PBS) at pH 7.4 (P4417, Merck) at a concentration of 0.6 mg/mL according to the manufacturer’s instructions. For labelling, the pH of 50 µg of protein solution was adjusted to pH 8.3 using 1 M NaHCO_3_ followed by addition of Alexa Fluor™ 647 NHS ester at a molar dye-to-protein ratio of 3:1. Samples were incubated at 4 °C overnight, and free dye was removed by size-exclusion chromatography on an ÄKTA pure system (Cytiva) equipped with a Superdex 200 Increase 3.2/300 column (Cytiva) using PBS at pH 7.4 as a buffer. Labeled and purified proteins were stored at −80 °C in PBS pH 7.4 containing 10% (w/v) glycerol as cryoprotectant.

### Affinity measurements of binary equilibrium by microfluidic diffusional sizing

To determine the affinity of ACE2–SARS-CoV-2 spike S1 binding, Alexa Fluor™ 647 labeled ACE2 (10 nM) was mixed with unlabeled SARS-CoV-2 spike S1 at increasing concentrations (100 pM–3.2 µM) in the presence of 90% heat-inactivated human serum (H5667, Merck) and incubated at 4 °C overnight. To determine the affinity of a neutralizing antibody to SARS-CoV-2 spike S1, fluorescently labeled SARS-CoV-2 spike S1 (10 nM) was mixed with unlabeled antibody (SAD-S35, ACROBiosystems) at increasing concentrations (3.8 pM–250 nM) in the presence of 90% heat-inactivated human serum (H5667, Merck) and incubated at 4 °C overnight. To measure complex formation by microfluidic diffusional sizing, 5 µL of sample were pipetted on a microfluidic chip and analysis was performed at the 1.5 nm–8.0 nm setting on a Fluidity One-W Serum (Fluidic Analytics). Serum autofluorescence was determined in the absence of labeled protein and used to correct MDS data measured of binding interactions. Error bars shown in figures are standard deviations from triplicate measurements. The equilibrium dissociation constant (*K*_D_) was determined by nonlinear least-squares (NLSQ) fitting (Prism, GraphPad Software) in terms of the following binary equilibrium:

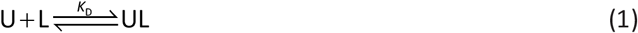

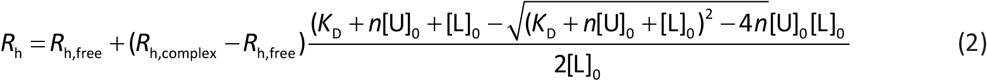

with *R*_h_, *R*_h,free_, and *R*_h,complex_ being the effective hydrodynamic radii at equilibrium, of the unbound labeled species and of the complex of unlabeled and labeled species, respectively. The parameter *n* is equivalent to the number of its binding sites. Furthermore, [U]_0_ and [L]_0_ are total concentrations of unlabeled and labeled species, respectively.

### Measurement of ternary equilibrium (recombinant proteins) by microfluidic diffusional sizing

To assess inhibition of ACE2–S1 binding by a recombinant NAb, a mixture of Alexa Fluor™ 647 labeled ACE2 and unlabeled S1 at concentrations of 10 nM and 50 nM, respectively, was equilibrated with increasing concentrations of NAb (30 pM–1.0 µM) in the presence of 90% heat-inactivated human serum (H5667, Merck) at a temperature of 4 °C overnight. MDS data was obtained as described above.

The following ternary equilibrium model^23^ was used for data analysis:

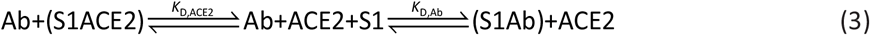

In this ternary mixture, the two *K*_D_ values are defined as follows:

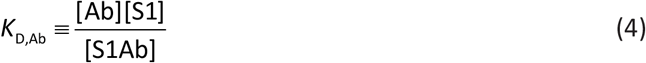

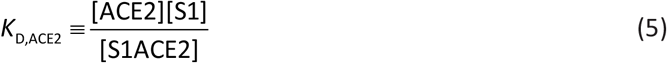

[Ab], [S1], [ACE2], [S1Ab], and [S1ACE2] are equilibrium concentrations of antibody, S1, ACE2, S1–antibody complex, and S1–ACE2 complex, respectively.

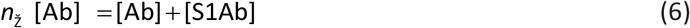

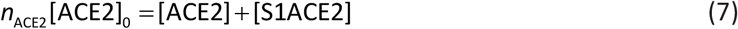

*n* can be used to determine binding stoichiometry.

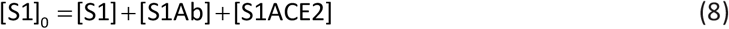

Substitution of Equations 6 and 7 into Equations 4 and 5, respectively, yields:

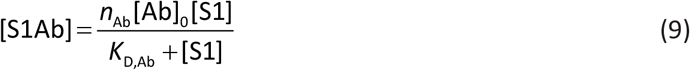

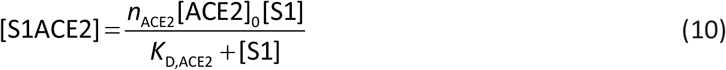

Insertion of Equations 9 and 10 into Equation 8 leads to:

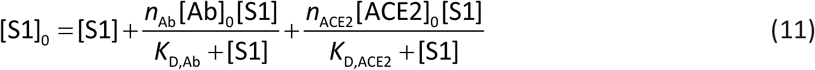

Rearrangement gives a cubic equation:

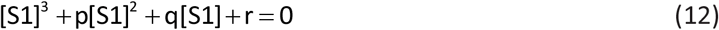

The coefficients are:

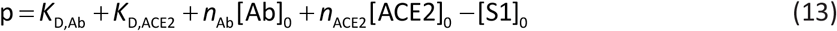

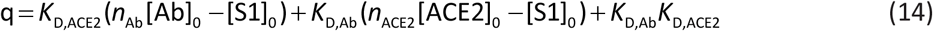

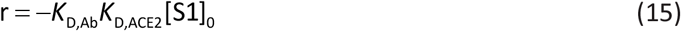

The following root is the only physically meaningful solution:

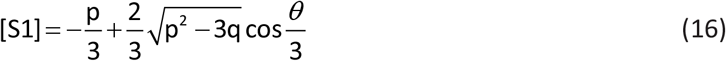

in which

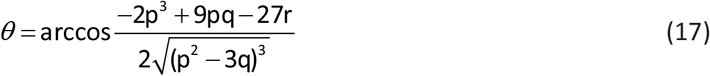

Equation 16 describes the equilibrium concentration of unbound S1 using the total concentrations of the individual components, two *K*_D_ values, and *n*.

### Global analysis of binary and ternary binding experiments

To extract equilibrium concentrations and corresponding bound fractions of ACE2 and S1 in the ternary mixture with NAb, we globally fitted (NLSQ) binary (ACE2–S1 and S1–NAb) and ternary (ACE2–S1–NAb) experiments. The binary experiments were fitted in terms of Equation 2, while the ternary experiment was fitted in terms of Equations 13–17. *K*_D,ACE2_, *K*_D,NAb_, and *R*_h,freeACE2_ were set to be global fitting parameters, while all other parameters were fitted locally. The stoichiometry *n* was set to 1 as concentrations refer to binding sites.

### Origin of serum samples

Anti-SARS-CoV-2 seropositive human serum samples (convalescent) were obtained from BioIVT. BioIVT sought informed consent from each subject, or the subjects legally authorized representative and appropriately documented this in writing. All samples are collected under IRB-approved protocols.

### Microfluidic antibody-affinity profiling of serum antibodies (MAAP)

MAAP was used to determine concentration of antibody-binding sites, [Ab], and *K*_D_ of antibodies in serum samples of SARS-CoV-2 seropositive individuals. To do so, fluorescently labeled SARS-CoV-2 spike S1 was mixed at constant concentrations of 10 nM, 40 nM, and 100 nM with serum at increasing concentrations (1%–90%) to generate three equilibrium binding isotherms per serum sample. MDS data was obtained as described above, the only difference being that samples were incubated for 1 h at 4 °C before measurement. Autofluorescence of serum samples was determined in the absence of labeled protein and used to correct MDS data measured of binding interactions. To determine the concentration of antibody binding sites and *K*_D_, global NLSQ fitting in terms of Equation 2 was used, having *R*_h,free_, *R*_h,complex_, [U]_0_ = [Ab] and *K*_D_ as global fit parameters.

### Measurement of ACE2–S1 binding inhibition by serum antibodies using microfluidic sizing

For sample 1, Alexa Fluor™ 647 labeled ACE2 and unlabeled S1 at concentrations of 10 nM and 30 nM, respectively, were mixed with increasing concentrations of serum and incubated at 4 °C for 1 hour. For sample 2, Alexa Fluor™ 647 labeled ACE2 and unlabeled S1 at concentrations of 5 nM and 20 nM, respectively, were mixed with increasing concentrations of serum and incubated at 4 °C for 1 hour. MDS data was obtained as described above.

## Supplementary Information

**Table S1.** Summary of thermodynamic parameters and 95% confidence intervals obtained for antibody-mediated inhibition of ACE2–S1 binding by individually fitting binary equilibria or by globally fitting binary and ternary equilibria.

| parameter | type of fit | best fit | 95% CI | Figure |
| --- | --- | --- | --- | --- |
| ACE2-S1 in 90% serum |  |  |  |  |
| $K_{D,ACE2}$ (nM) | binary | 7.0 | (3.5–12.6) | 1B |
| antibody affinity in serum sample 1 |  |  |  |  |
| $K_{D,Ab}$ (nM) | binary | 0.7 | (0.01–2.1) | S1 |
| $K_{D,Ab}$ (nM) | global | 0.8 | (0.1–2.4) | 3A |
| antibody affinity in serum sample 2 |  |  |  |  |
| $K_{D,Ab}$ (nM) | binary | <0.7 | (n.d.–0.7) | S1 |
| $K_{D,Ab}$ (nM) | global | <0.5 | (n.d.–0.5) | 3C |

**Figure S1.**
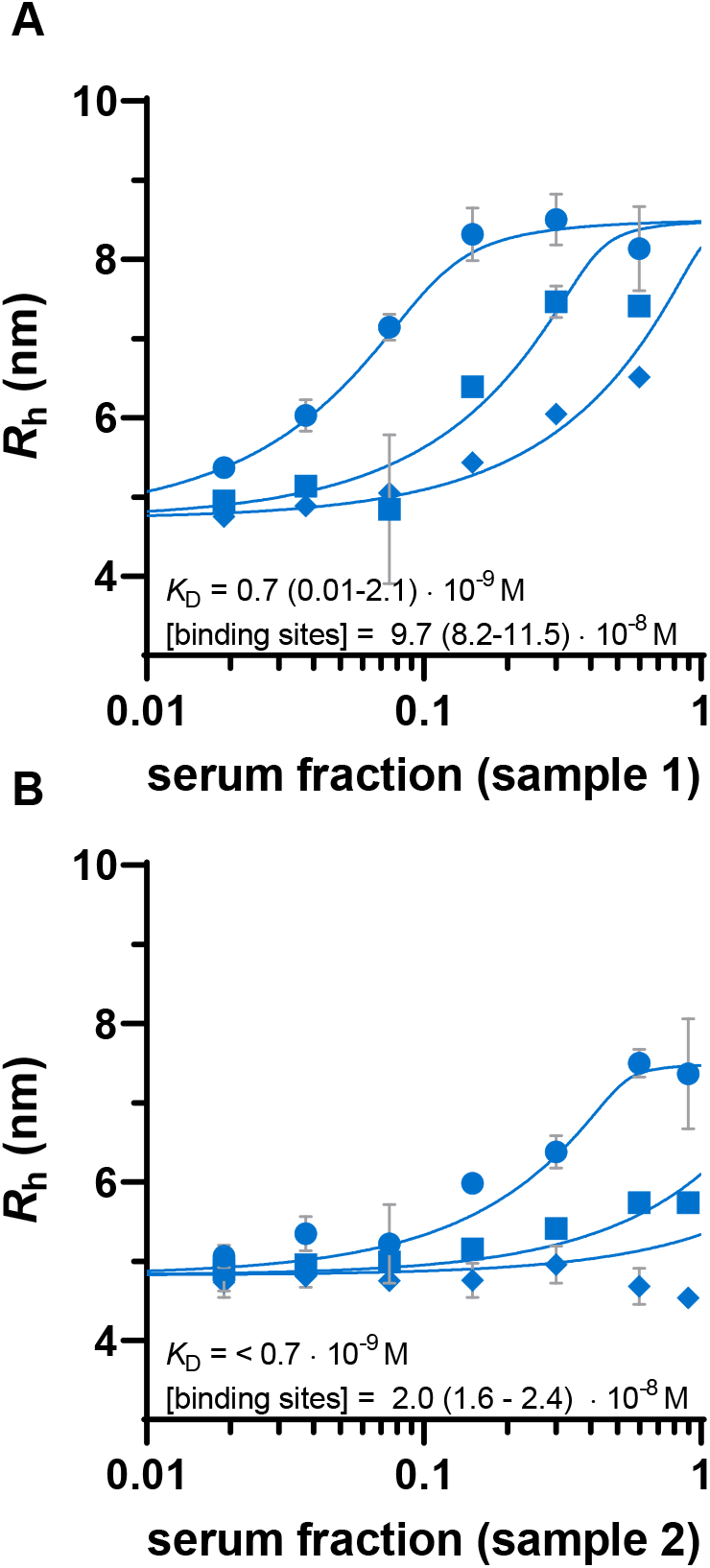
Microfluidic antibody-affinity profiling in serum obtained from two anti-SARS-CoV-2 sero-positive individuals. A) and B) Fluorescently labeled S1 at concentrations of 10 nM (circles), 40 nM (squares) or 100 nM (diamonds) was mixed with increasing concentrations of serum, and binding of antibodies was assessed as an increase in *R*_h_. *K*_D_ and concentration of antibodies were obtained by NLSQ in terms of **Equation 2**, having *K*_D_, [U]_0_ (antibody concentration), and *R*_h_ of the complex as global fitting parameters. Error bars are standard deviations obtained from duplicate measurements.

## Notes

### Competing Interest Statement

TPJK is a member of the board of directors of Fluidic Analytics. AA is a member of the scientific advisory committee of Fluidic Analytics. SF, MAP, VD, ASM, AI, AYM, SRAD, and HF are employees of Fluidic Analytics. All other authors declare no competing interests.

## References

1. WHO. Coronavirus disease (COVID-19) Weekly Epidemiological Update Global epidemiological situation. (2020).

2. Wajnberg, A. et al.. SARS-CoV-2 infection induces robust, neutralizing antibody responses that are stable for at least three months. medRxiv (2020) doi:10.1101/2020.07.14.20151126.

3. Garritsen, A. et al.. Two-tiered SARS-CoV-2 seroconversion screening in the Netherlands and stability of nucleocapsid, spike protein domain 1 and neutralizing antibodies. medRxiv (2020) doi:10.1101/2020.10.07.20187641.

4. Emmenegger, M. et al.. Early peak and rapid decline of SARS-CoV-2 seroprevalence in a Swiss metropolitan region. medRxiv 22 (2020) doi:10.1101/2020.05.31.20118554.

5. Brouwer, P. J. M. et al.. Potent neutralizing antibodies from COVID-19 patients define multiple targets of vulnerability. Science 369, 643–650 (2020).

6. Cao, Y. et al.. Potent Neutralizing Antibodies against SARS-CoV-2 Identified by High-Throughput Single-Cell Sequencing of Convalescent Patients’ B Cells. Cell 182, 73–84 (2020).

7. Ju, B. et al.. Human neutralizing antibodies elicited by SARS-CoV-2 infection Plasma antibody response against SARS-CoV-2. Nature 584, 115–119 (2020).

8. Liu, L. et al.. Potent neutralizing antibodies against multiple epitopes on SARS-CoV-2 spike. Nature 584, 450–456 (2020).

9. Rogers, T. F. et al.. Isolation of potent SARS-CoV-2 neutralizing antibodies and protection from disease in a small animal model. Science 369, 956–963 (2020).

10. Seydoux, E. et al.. Analysis of a SARS-CoV-2-Infected Individual Reveals Development of Potent Neutralizing Antibodies with Limited Somatic Mutation. Immunity 53, 98–105 (2020).

11. Wec, A. Z. et al.. Broad neutralization of SARS-related viruses by human monoclonal antibodies. Science 369, 731–736 (2020).

12. Wu, Y. et al.. A noncompeting pair of human neutralizing antibodies block COVID-19 virus binding to its receptor ACE2. Science 368, 1274–1278 (2020).

13. Zost, S. J. et al.. Potently neutralizing and protective human antibodies against SARS-CoV-2. Nature 584, 443–449 (2020).

14. Piechotta, V. et al.. Convalescent plasma or hyperimmune immunoglobulin for people with COVID-19: a living systematic review. Cochrane Database Syst. Rev. (2020) doi:10.1002/14651858.CD013600.pub2.

15. Burton, D. R. & Walker, L. M. Rational Vaccine Design in the Time of COVID-19. Cell Host Microbe 27, 695–698 (2020).

16. Hoffmann, M. et al.. SARS-CoV-2 Cell Entry Depends on ACE2 and TMPRSS2 and Is Blocked by a Clinically Proven Protease Inhibitor. Cell 181, 271–280 (2020).

17. Nie, J. et al.. Establishment and validation of a pseudovirus neutralization assay for SARS-CoV-2. Emerg. Microbes Infect. 9, 680–686 (2020).

18. Oguntuyo, K. Y. et al.. Quantifying absolute neutralization titers against SARS-CoV-2 by a standardized virus neutralization assay allows for cross-cohort comparisons of COVID-19 sera. medRxiv (2020) doi:10.1101/2020.08.13.20157222.

19. Tan, C. W. et al.. A SARS-CoV-2 surrogate virus neutralization test based on antibody-mediated blockage of ACE2–spike protein–protein interaction. Nat. Biotechnol. 38, 1073–1078 (2020).

20. Zhang, L. et al.. Anti-SARS-CoV-2 virus antibody levels in convalescent plasma of six donors who have recovered from COVID-19. Aging (Albany. NY). 12, 6536–6542 (2020).

21. Schneider, M. M. et al.. Microfluidic Affinity Profiling reveals a Broad Range of Target Affinities for Anti-SARS-CoV-2 Antibodies in Plasma of Covid Survivors. medRxiv (2020) doi:10.1101/2020.09.20.20196907.

22. Schneider, M. M. et al.. Microfluidic Antibody Affinity Profiling for In-Solution Characterisation of Alloantibody - HLA Interactions in Human Serum. bioRxiv (2020) doi:10.1101/2020.09.14.296442.

23. Krainer, G. & Keller, S. Single-experiment displacement assay for quantifying high-affinity binding by isothermal titration calorimetry. Methods 76, 116–123 (2015).

